# Inhibition of Proliferation of Acute Lymphocytic Leukemia (ALL) by frequency-specific Oscillating Pulsed Electric Fields (OPEF) broadcast by an enclosed gas plasma antenna

**DOI:** 10.1101/2023.05.03.539248

**Authors:** Anthony Holland, James Bare

## Abstract

Frequency-specific Oscillating Pulsed Electric Fields (OPEF) broadcast by an enclosed gas plasma antenna with a frequency of 160kHz from a distance of 18 inches inhibited the proliferation of Acute Lymphocytic Leukemia cells in vitro by up to 44%.

## INTRODUCTION

Acute Lymphocytic Leukemia (ALL) affects some 6,600 people in the United States annually and about 1,560 die of it per year with 80 % of those deaths being adults ^(1)^. It is the most common malignancy of childhood, primarily occurring in children between the ages of two and ten ^(2)^.

Medical treatment protocols for ALL are initially based upon a multidrug chemotherapeutic regimen and should relapse occur, CAR-T and bone marrow transplants are often used ^(3)^. All treatment methods of ALL have potentially serious and occasionally fatal side effects. Many patients, especially children, show a strong initial response to treatment with 90+ % of children going into remission, and for adults, 60 to 80% reach remission. Adult patients that relapse after their initial remission have an approximate one-year survival, even if they achieve a second complete remission ^(3)^. Almost 600 children a year who initially achieved remission, relapse.

Currently only about 30 to 50% of these children will survive their first relapse. Of those that again obtain a second remission, they too may relapse at a later date. With each relapse the chances of the child being cured decreases ^(4)^.

In spite of advances in treatment, a large number of patients that relapse become refractory to further treatment. Relapse treatment involves the utilization of aggressive high dosage chemotherapy, and at times, stem cell transplants. The overall expense of treatment for relapse can be daunting. A stem cell transplant costs over $350,000 to $800,000 ^(5)^. Given the yearly number of deaths following relapse, a potential new method of ALL treatment is evaluated, a method which as a potential future medical treatment is nontoxic to healthy cells ^(6)^, noninvasive, and potentially inexpensive. Frequency-Specific Oscillating Pulsed Electric Fields (OPEF) broadcast by an enclosed gas plasma antenna were experimentally tested to determine effects upon ALL cell growth and proliferation. Oscillating Pulsed Electric Fields (OPEF) rely upon pulse rates that are specific to cell size and type ^(7, 8)^ but the fields do not vary in their rate of oscillation (27.12Mhz), only the pulse repetition rate (PRR) is varied to create a specific frequency of the pulses.

Intermediate frequency alternating current (oscillating, but non-pulsed) electric fields between 100kHz and 500kHz and of low intensity (1-3V/cm), sometimes known as “tumor treating fields” (TTF), are now an accepted FDA approved cancer treatment therapy in world-wide use ^(9)^. TT fields have shown clinically proven efficacy to inhibit the growth of various cancers both in vitro and in vivo and to induce apoptosis and necrosis in numerous different cancer cell lines ^(6)^. TT field therapies have FDA approvals for the treatment of brain gliomas and mesothelioma ^(10,11)^. TT fields are undergoing clinical trials for treatment of Pancreatic Cancer, Non-Small Cell Lung Cancer, Ovarian Cancer, Liver Cancer and Gastric Cancer ^(12)^. Electric Field Frequencies unique to specific cancer cell types have been shown to be nontoxic to quiescent healthy cells and demonstrate inhibition of proliferation of fast-growing cancer cells ^(6)^.

A substantial body of peer-reviewed literature has established the science behind the use of frequency-specific electric fields in the treatment of cancer ^(13)^. One company, Novocure, is making notable progress through numerous clinical trials and for many different types of cancer using what they have deemed as Tumor Treating Fields (TTF) ^(12)^. However, to date, the FDA approved medical treatments of frequency-specific electric fields have been confined to a method that utilizes specially insulated transducers attached to the outside of the body and placed at pre-determined angles to a specific location of a known tumor ^(14)^. Treatment of cancer cells using alternating electrical fields requires a certain field strength to obtain effectiveness. As there is a limit as to how much power (volts and current) can be applied through one electrode, Novocure uses an array of electrodes that are focused upon a particular tumor to obtain the necessary field strength ^(14, 15)^. While this method has been demonstrated to be very effective, this approach does not allow for the treatment of systemic cancers, which have no one specific location in the body or are widely spread throughout the body, as in leukemia or metastatic cancers. Such systemwide cancers require a new method of delivery of the frequency-specific electric fields, a method which could potentially treat the entire body at once by using a projected and pulsed electric field of a specific PRR frequency emitted from an enclosed gas plasma antenna that inherently creates strong electric fields ^(16)^.

OPEF fields transmitted by an enclosed gas plasma antenna can easily overcome the power limitations of electrodes. The field may easily be increased as might be required to provide sufficient strength to act upon the whole body at depth. OPEF electric field strength can be increased in three manners: increase in the driving power of the plasma tube, increase in the number of gas molecules within the plasma tube (larger tube/higher gas pressures), and closeness of the plasma tube to the tumor ^(16)^. In contrast to localized TT field treatment, OPEF offers the potential to globally treat metastatic cancers, and cancers of the blood or lymphatic system.

While the experiments described in this document refer to an effective treatment of cells in vitro from a distance of 18 inches (45.72 cm), our previous experiments have shown substantial reduction in cancer cell numbers from a distance of up to 24 inches (60 cm) ^(17)^. TT fields rely upon mitotic spindle disruption that results in improper chromosome segregation and mitotic catastrophe ^(18)^. OPEF’s transmitted via an enclosed gas plasma antenna have the ability to replicate the effects of TT fields, but we have also found that different frequency ranges (PRR) can have unique detrimental effects upon cancer cells. Notable PRR frequency-specific responses of cancer cells include the induction of surface tension in the actomyosin cortical layer of mitotic cells which can potentially cause the cells to burst ^(19)^.

## METHODS

Frequency-specific Oscillating Pulsed Electric Fields (OPEF) projected by an enclosed gas plasma antenna were tested against Acute Lymphocytic Leukemia cells (ALL) in vitro (Coriell Institute Catalog #GM03638) maintained in RPMI 1640 cell media (Life Technologies Catalog #61870036) supplemented with 20% fetal bovine serum (Life Technologies catalog #A3160601). Flasks were seeded at 320,000 cells/mL and incubated in a humidified 5% CO2 incubator at 37 degrees C for a period of 24 hours following initial seeding. Flasks were treated with the electronic OPEF signals according to the following schedule: 12-3pm cells on heated (37 degrees C) microscope stage treated with OPEF, 3-4pm cells returned to incubator, 4-6pm cells treated with OPEF on heated scope stage, 6-7pm cells returned to incubator, 7-9pm cells treated with OPEF on heated scope stage, 9-10pm cells returned to incubator, 10pm-12pm midnight cells treated with OPEF on heated microscope stage. This protocol resulted in a total of 9 hours of OPEF treatment daily, followed by 12 hours in incubator. This procedure was repeated for days two and three. After three days of the described daily treatment protocol, cells were incubated an additional 48 hours and then assayed using the Trypan Blue exclusion method. Control experiments were run prior to the start of OPEF treatments and followed the exact same protocols as described above but without OPEF treatment. For the five OPEF treatment experiments, a PRR of 160kHz was chosen based on previous heuristic experiments (unpublished).

## ELECTRONICS SETUP

OPEF signals were generated with the following electronic equipment: Tenma 72-7655 DC Power Supply, F125 Square Wave Generator (Atelier Robin), duty cycle 50%, OM2 custom RF transmitter (Plasmasonics) ^(20)^ operating with a carrier frequency of 27.12 MHz, 300 Watt PEP Class C custom RF amplifier (Plasmasonics) that typically provides about 1000 volts to the Phanotron plasma tube, MFJ customized Antenna Impedance Tuner (Plasmasonics), and a Large Phanotron plasma antenna tube (Bill Cheb, billplasmatubes.com). The Phanotron tube is evacuated and backfilled with helium and placed 18 inches (45.72 cm) from the center of the heated microscope stage where the leukemia cells were placed for treatment.

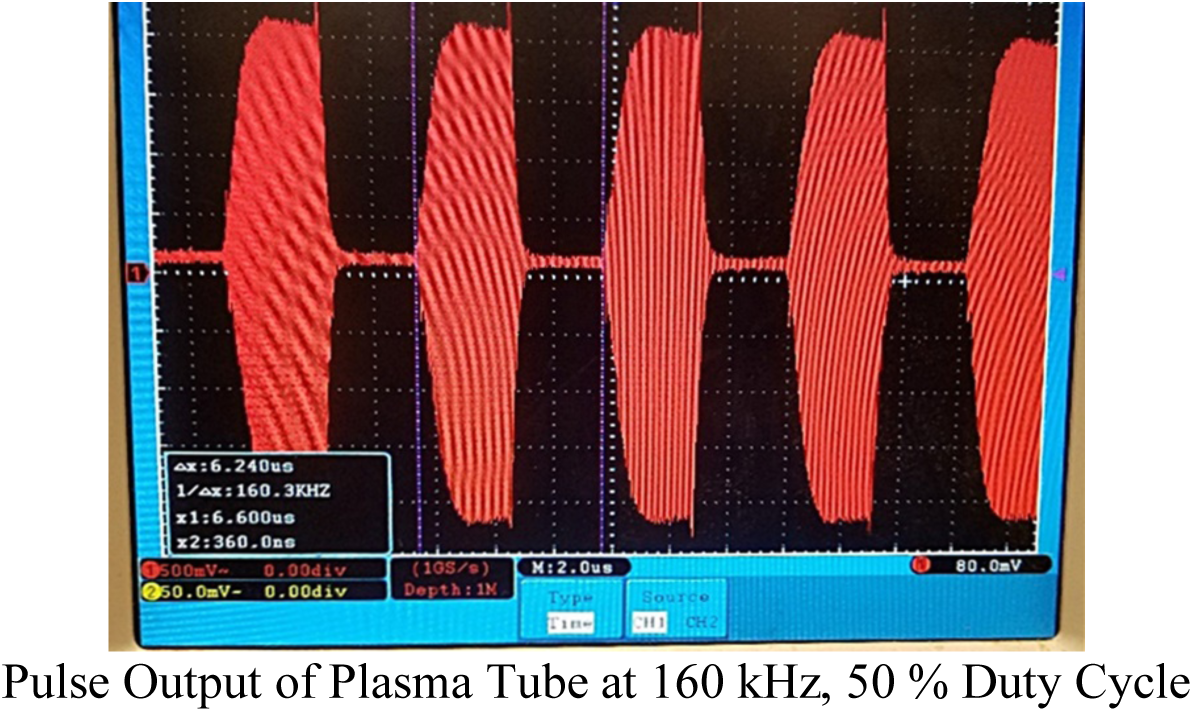

The 160 kHz square wave signal was pulse amplitude modulated by the OM-2. The OM-2 is a patented RF pulse transmitter that creates pulses via overmodulation (an index of modulation exceeding 1). The signal is pulsed at a rate that equals the fundamental frequency of the square wave. Because the signal is pulsed, the biological effects seen against cancer cells are non-thermal in nature ^(21)^. The pulse repetition rate (PRR) works in a synergistic relationship with the strength of the electric field. The strength of the electric field has been calculated in our previous publications ^(7)^.

## RESULTS

The average growth rate of five control experiments was 3.132. When compared to the average growth rate of the five control experiments, OPEF treated cells showed an inhibition of proliferation with growth rate reductions of 42%, 43%, 37%, 24% and 44% respectively. The average growth rate reduction in the five OPEF experiments was 38% (P = 0.0053).

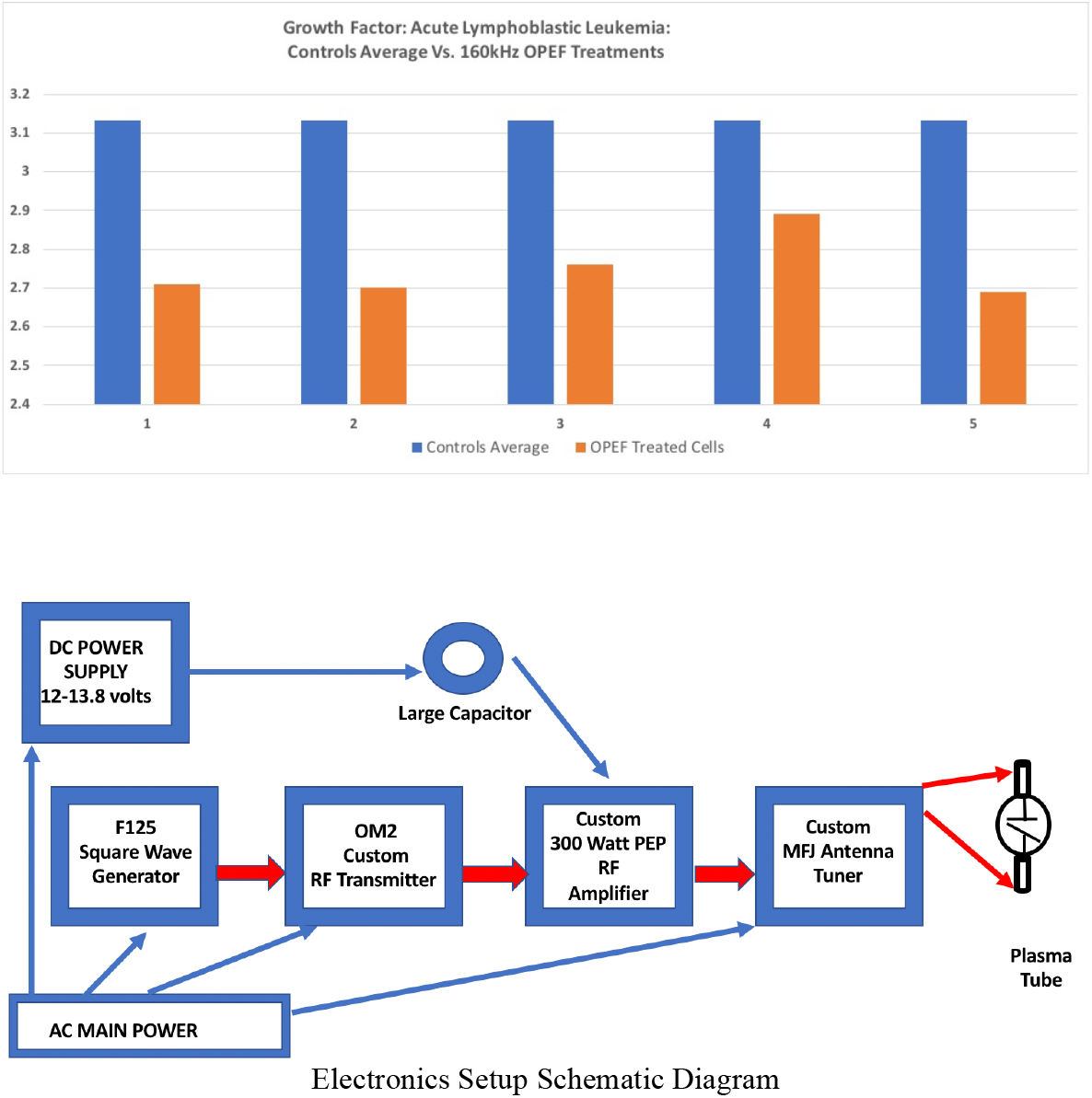

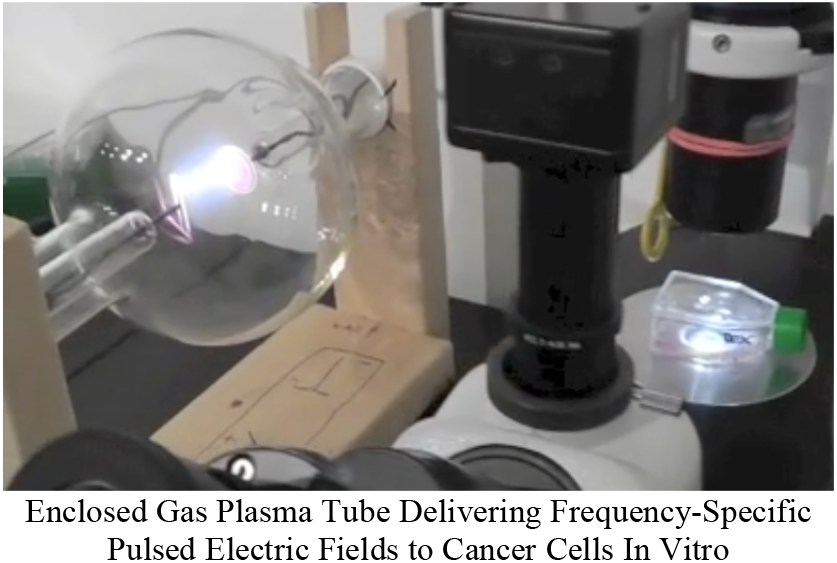

## DISCUSSION

The majority of OPEF treatment experiments resulted in an average growth rate reduction of 43%. These results are in line with the inhibition of proliferation of cancer cell growth seen in our other experiments that utilize frequency-specific electric fields ^(17)^, but a unique aspect of our experiments is that the frequency-specific electric fields are pulsed and are ‘broadcast’ by an enclosed gas plasma antenna located 18 inches from the cancer cells. These proof of concept experiments suggest that this method could eventually become viable as a future cancer treatment that treats the entire body of a patient at once, thus addressing systemic cancers Of the blood, lymphatic system or metastatic cancers in general.

Thanks to Professor Tom O’Connell, Skidmore College, for assistance in computing the data P value and to Frederic Bellossi (E.E.), for special assistance.

